# Alteration of fecal microbial compositions and bacterial taxa in female osteoporotic patients

**DOI:** 10.1101/2020.01.21.914903

**Authors:** Cuifeng Zhu, Jianguo Liu, Yong Pan, Ye Fan, Leigang Jin, Yan Liu, Zhentian Zhang, Yu Gan, Wei Tang, Jinhuan Li, Zhuang Xiong, Genming Xu, Xiuping Lin, Yuan Zhang, JinChuan Cai, Muxiu Yang, Leijun Zhang, Liehui Xiao, Yi Pan, Kejian Wang, Aimin Xu

**Author notes:** Leading corresponding author: Aimin Xu, Professor, University of Hong Kong & Director of the State Key laboratory of Pharmaceutical Biotechnology.; Contribution to the study: professor, director, responsible for the overall design and guidance of the project). Co-corresponding author: Kejian Wang, Associate Professor of Guangzhou Medical University, (Contributions to this study: data analysis and processing of intestinal flora structure and gene polymorphism detection, and guidance of paper writing) and Yi Pan, Associate Professor of Hunan Engineering Research Center for Obesity and Its Metabolic Complications, Xiangya Hospital, Central South University,. (Contributions to this study: post doctorate, Responsible for the establishment, detection, analysis and quality control of the database of intestinal flora structure and gene polymorphism). Cuifeng Zhu, Jianguo Liu and Yong Pan contributed equally to this work. Author order was determined on the basis of seniority. (CuiFeng Zhu, Department director of Dept of Clinical Nutrition, chief physician, master tutor of Shenzhen hospital of southern medical university. Post doctor fellows of Dept of endocrinology of university of Hong Kong & The State Key laboratory of Pharmaceutical Biotechnology. (Contributions to this study: chief physician of Nutrition department and post doctorate, responsible for the design and implementation of this project, data collation and paper writing); JianGuo Liu, Associate chief physician, Department director of Dept of Health management of Shenzhen hospital of southern medical University. (Contribution to this study: general practitioner at health management center, responsible for case collection and data collation); Pan Yong, Post doctor fellows of Dept of endocrinology of university of Hong Kong & The State Key laboratory of Pharmaceutical Biotechnology. (Contributions to this study: post doctorate, responsible for data collation and thesis writing guidance of this project). Other author: Ye Fan: School of Stomatology and Medicine, Foshan University, Foshan, China, 528000. Tele (Contribution to this study: visiting scholar, responsible for the data processing of intestinal microflora gene polymorphism test results). Leigang Jin: Department of Medicine, Li Ka Shing Faculty of Medicine, University of Hong Kong & The State Key laboratory of Pharmaceutical Biotechnology, Hong Kong, China. (Contributions to this study:doctor student, responsible for the early data processing of intestinal microflora gene polymorphism). Yan Liu: Shenzhen Hospital of Southern Medical University, Shenzhen, Guangdong province, 518000, 13570469574 (Contribution to this study:Nutrition department attending physician, in charge of the medical history of selected patients, menstrual history, life history and other data survey). Zhentian Zhang: Shenzhen Hospital of Southern Medical University, Shenzhen, Guangdong province, 518000., 13544154119. (Contribution to this study: a nutritionist was responsible for the investigation of intestinal mucosal barrier function in selected patients with medical history and other data). Yu Gan: The Third Affiliatd Hospital of Guangzhou Medical University,. (Contribution to this study: biomedical engineer, responsible for the analysis and processing of intestinal flora structure and gene polymorphism detection data). Wei Tang: Yearth Biotechnology Co. Ltd., Changsha, China, 410000, 0731-88709258. (Contribution to this study:biomedical engineer, responsible for the extraction of fecal intestinal flora RNA and reverse transcription of DNA). Jinhuan Li: Yearth Biotechnology Co. Ltd., Changsha, China, 410000,. 0731-88709258. (Contribution to this study:biomedical engineer, responsible for the extraction of fecal intestinal flora RNA and reverse transcription of DNA). Zhuang Xiong: Yearth Biotechnology Co. Ltd., Changsha, China, 410000,. 0731-88709258. (Contribution to this study: biomedical engineer, responsible for the detection and analysis report of fecal intestinal flora gene polymorphism). Genming Xu: Yearth Biotechnology Co. Ltd., Changsha, China, 410000,. 0731-88709258. (Contribution to this study: biomedical engineer, responsible for the detection and analysis report of fecal intestinal flora gene polymorphism). Xiuping Lin: Shenzhen Hospital of Southern Medical University, Shenzhen, Guangdong province, 518000., 13500053960. (Contribution to this study: head nurse at health management center, responsible for case collection and collection of serum and stool samples from perimenopausal women). Yuan Zhang: Shenzhen Hospital of Southern Medical University, Shenzhen, Guangdong province, 518000., 15118092923. (Contributions to this study: Health management Centre general practitioner, perimenopausal female case collection and collection of serum and stool samples). JinChuan Cai: Shenzhen Hospital of Southern Medical University, Shenzhen, Guangdong province, 518000. E-mail:, 13600158985. (Contributions to this study: Technician at the health management center, bone density measurement.). Muxiu Yang: Shenzhen Hospital of Southern Medical University, Shenzhen, Guangdong province, 518000., 13823236331. (Contributions to this study:: Biomedical engineer, responsible for bone density analysis in this study). Leijun Zhang: Shenzhen Hospital of Southern Medical University, Shenzhen, Guangdong province, 518000., 13802278833. (Contributions to this study:Orthopedic surgeon at health management center, case collection). Liehui Xiao: Shenzhen Hospital of Southern Medical University, Shenzhen, Guangdong province, 518000., 13189097298. Contributions to this study: integrated medicine physician, case collection). Author Contribution:Cuifeng Zhu1,2,3,★, Jianguo Liu 1★, Yong Pan2,3★, Yi Pan4, Ye Fan5,Leigang Jin2,3, Yan Liu1, Zhentian Zhang1, Yu Gan6, Wei Tang7, Jinhuan Li7, Zhuang Xiong7, Genming Xu7, Xiuping Lin1, JinChuan Cai1,Yuan Zhang1, Muxiu Yang1, Leijun Zhang1, Liehui Xiao1, Kejian Wang6#, Aimin Xu2,3.

## Abstract

**Background:** Gut microbiota, mainly characterized by fecal bacterial compositions, affects human immune system and pathophysiological development. Our aim was to measure the quantitative differences of fecal bacterial compositions between osteoporotic patients and healthy subjects, and to identify novel bacterial taxa that speculate the incidence of osteoporosis in female.

**Method:** We recruited 104 female subjects, including 45 osteoporotic individuals and 59 healthy control. Fecal samples were collected for further analysis by 16S rRNA quantitative arrays and bioinformatics analysis.

**Results:** Analyses of α- and β-diversity demonstrated that the diversity and composition of fecal bacterial compositions were both significant different in osteoporosis group, as compared with healthy group. Multiple bacterial genera were significantly increased (e.g., Roseburia and Bacteroides) or decreased (e.g., Streptococcus and Dorea) in the osteoporotic cases. Furthermore, the osteoporosiscould be efficiently determined by the random forest model based on differential taxa (area under ROC curve = 0.93).

**Conclusion:** There were obvious different fecal microbial characteristics between female osteoporosis and healthy subjects. These findings provided evidence for understanding the host-gut microbiota interplay in female osteoporosis, and supported clinical applications of gut microbiota analysis for female osteoporosis diagnosis

## Introduction

Osteoporosis is characterized by low bone mass and increased bone fragility (1, 2). The deterioration of bone tissue architecture in osteoporosis is mainly caused by imbalanced bone formation slower than bone resorption (3), which increases the incidence of bone fractures. Osteoporotic fractures result in a heavy burden to health services worldwide, especially increased morbidity and reduced survival in the elderly(4, 5).

Conventional therapeutic interventions on osteoporosis include general lifestyle changing (e.g., balanced diet and regular exercise) (6), and intake of approved drugs (e.g., oestrogen and bisphosphonates) (7). In recent years, a series of studies suggested that gut microbiota colonizing in the intestinal tract not only affected the nutrition metabolism, but also contributed to the occurrence and progression of osteoporosis (8). Das et al. reported there were differences of gut microbiota composition between osteoporosis /osteopenia and controls with an Irish cohort of older adults (9). Wang et al. analyzed the gut microbiota diversity in a relatively small sample with 6 primary osteoporosis patients, 6 osteopenia patients and 6 healthy controls (10). Li et al. identified several differential fecal bacterial taxa from patients with low bone mineral density (11). However, the above research findings were based on subjects without considering sex, and the results were also inconsistent.

Osteoporosis is a typical disease of females. One key reason is the sudden decline of estrogen production for women after the menopause but a gradual process for men (12). Therefore, hormone replacement therapy effectively decreased the risk of bone fracture in post-menopausal women (13, 14). It indicated that the pathophysiology of osteoporosis may differ between women and men. However, current fecal microbial studies on osteoporosis mixed female and male subjects together for analysis, leading to the misunderstanding of female-specific characteristics. To this end, the present study aimed to investigate the alterations in gut microbiota of female osteoporotic patients using 16S rRNA high-throughput sequencing analysis. The results could provide bioinformatics characteristics of dysbiosis in osteoporoticwomen, thus helping to guide probiotics treatment and primary prevention.

## Materials and Methods

### Patient recruitment and bone mineral density examination

This study was approved by the Ethics Committee of Shenzhen Hospital of Southern Medical University. Written informed consent was signed by all participants prior to enrollment. BMD values at skeleton sites of the lumbar spine and the total hip joint were obtained using dual-energy X-ray absorptiometry (DXA) (DiscoveryTM CI, QDR SERIES, HOLDGIC). Subject with BMD T-Score < −2.5 were classified into osteoporotic group, whereas participants with BMD T-Score > 1.0 were classified into the control group. Those subjects meeting the following criteria were excluded for study: (1) use of antibiotics or hormone within 6 months before enrollment, (2) undergoing gastrointestinal diseases, and (3) history of hypothyroidism or hyperthyroidism. Finally, 104 participants (45 osteoporosis cases and 59 controls) were included in analysis.

### Fecal sample collection and DNA extraction

Fresh fecal samples were collected in sterile tubes and frozen at −80 °C for further use. Genomic DNA was extracted from each fecal sample using QIAamp Fast DNA Stool Mini Kit (QIAGEN, Germany) according to the manufacturer’s instructions. The amount of DNA was determined by NanoDrop 2000 UV-via spectrophotometer (Thermo Scientific, USA). Integrity and size of DNA were checked by 0.8% (w/v) agarose gel electrophoresis in 0.5 mg/ml ethidium bromide.

### 16S rRNA sequencing

We amplified the V3-V4 hypervariable regions of 16S rRNA gene using the following PCR primers: forward 5’-CCTACGGGRSGCAGCAG-3’ and reverse 5’-GGACTACVVGGGTATCTAATC-3’. PCR amplification was performed in a mixture containing 1.0 μl of forward primer, 1.0 μl of reverse primer and 6.0 μl of DNA. The reactions were started at 96 °C for 30 s, followed by 40 cycles of 96 °C for 15 s, 60 °C for 15 s, and 75 °C for 15 s, with a final extension step at 75 °C for 1 min. The concentration of DNA libraries was quantified using PicoGreen DNA Assay (Invitrogen, USA). Pooled DNA libraries were diluted to 10 pManddenatured in 0.2N NaOH, mixed withPhiX control library (Illumina Inc., USA), and paired-end sequenced (2 × 250 bp) on the Illumina MiSeq platform.

### Sequencing data analysis

The Quantitative Insights Into Microbial Ecology pipeline was employed to process the sequencing data (15). Paired-end reads were merged using PANDAseq, sequences were de-noised using USEARCH (ver. 8.0.1623), and chimera checked with UCHIME26. Operational Taxonomic Units (OTUs) were picked at 97% similarity and representative sequences were generated. Sequences were aligned with PyNAST using Greengenes database and taxonomy assigned to the lowest possible taxonomic level using the Ribosomal Database Project Classifier at a 80% bootstrap value threshold. OTUs found in above 50% samples were retained. The numbers of sequences were normalized for further analyses.

The ACE, Chao, Shannon and Simpson index were calculated to assess α-diversity. Principal coordinate analysis (PCoA) and non-metric multidimensional scaling (NMDS) based on Bray-Curtis distance of OTUs were performed to provide an overview of the between-group bacterial difference (i.e, β-diversity). The Pearson’s correlation coefficient was calculated to evaluate the association between bacteria abundance and bone mass density. Statistical analyses and data visualization were performed using R software package (version 3.5.1).

## Results

### Characteristics of study subjects

A total of 45 osteoporosis patients and 59 healthy controls were recruited according to the inclusion and exclusion criteria (Table 1). There was no statistical difference in body mass index (BMI), alcohol intake, tobacco used and history of fracture, renal disease or osteoarthritis disease between two groups (p > 0.05). The osteoporotic cases were generally older than the control subjects (average age 59.07 vs 54.59). The osteoporosis group had significantly lower T-score, Z-score and BMD than the control group. The values of 25-oh-VD were decreased in osteoporosis cases as compared to control individuals. Notably, we also found significant lower level of diamine oxidase and higher level of D-lactic acid and lipopolysaccharide (LPS) in osteoporosis group than in control group (P < 0.001), suggesting undermined intestinal permeability in osteoporotic cases.

**Table 1.** Characteristics of osteoporosis and control subjects.

|  | Osteoporosis (n=45) | Control (n=59) | P-value<sup>#</sup> |
| --- | --- | --- | --- |
| Age (mean±SD) | 59.07±7.41 | 54.59±6.61 | 0.0017* |
| BMI (mean±SD) | 22.98±2.85 | 23.52±3.61 | 0.41 |
| Drinking, n (%) | 12 (26.7) | 15 (25.4) | 1.00 |
| Smoking, n (%) | 7 (15.6) | 9 (15.3) | 1.00 |
| T-Score | -2.74±0.57 | 0.16±1.01 | 8.74×10<sup>-32</sup>* |
| LS T-score | -3.06±0.45 | -0.18±0.41 | 2.22×10<sup>-57</sup>* |
| LS Z-score | -1.30±0.58 | 1.69±0.18 | 4.05×10<sup>-61</sup>* |
| LS BMD (g/cm<sup>2</sup>) | 0.65±0.06 | 0.88±0.10 | 1.24×10<sup>-24</sup>* |
| Hip T-score | -2.72±0.44 | 0.13±0.36 | 3.82×10<sup>-60</sup>* |
| Hip Z-score | -1.41±0.24 | 1.36±0.17 | 2.94×10<sup>-87</sup>* |
| Hip BMD (g/cm<sup>2</sup>) | 0.55±0.03 | 0.83±0.05 | 2.74×10<sup>-59</sup>* |
| 25-oh-VD | 17.69±4.78 | 45.01±6.58 | 1.29×10<sup>-42</sup>* |
| Diamine oxidase (U/L) | 1.22±0.45 | 3.05±0.58 | 4.71×10<sup>-32</sup>* |
| D-Lactic Acid (mg/L) | 19.82±4.49 | 7.12±1.34 | 1.02×10<sup>-37</sup>* |
| LPS (U/L) | 3.42±0.76 | 0.98±0.42 | 2.00×10<sup>-38</sup>* |
| Fracture, n (%) | 0 (0) | 0 (0) | 1.00 |
| Renal Disease, n (%) | 1 (2.2) | 0 (0) | 0.43 |
| Osteoarthritis Disease, n (%) | 3 (6.7) | 0 (0) | 0.078 |

<sup>#</sup> P-values were calculated with Fisher's exact test for nominal variables and Student's t-test for continuous variables.
\* Statistically significant difference between two groups.

### Comparisons of gut microbial diversity

We performed α-diversity analysis and found significant difference between osteoporotic and control groups. The osteoporosis cases exhibited higher values of ACE, Chao, Shannon indices and lower value of Simpson index (P < 0.01, Figure 1), suggesting that the microbial diversity is reduced in osteoporosis status. In addition, the analysis of the β-diversity calculated on Bray-Curtis dissimilarity and unweighted UniFrac distance both showed that microbiota composition of osteoporosis patients clustered apart from that of healthy controls (PERMANOVA P < 0.01, Figure 2), indicating between-group difference in microbiota composition.

**Figure 1.**
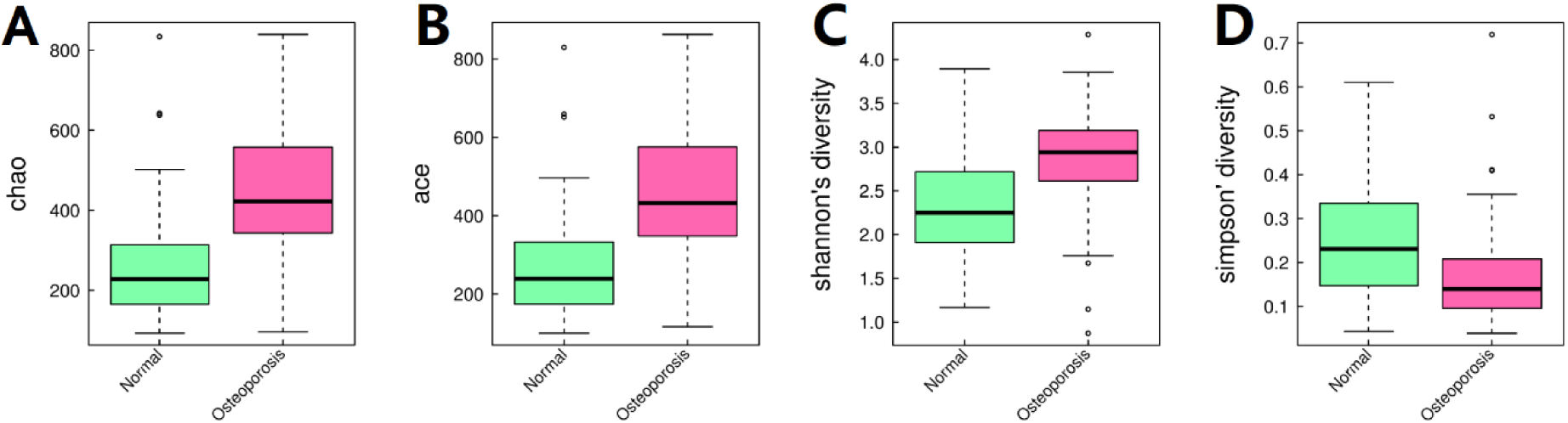
Comparison of α-diversity measured by ACE (A), Chao (B), Shannon (C) and Simpson (D) indices.

**Figure 2.**
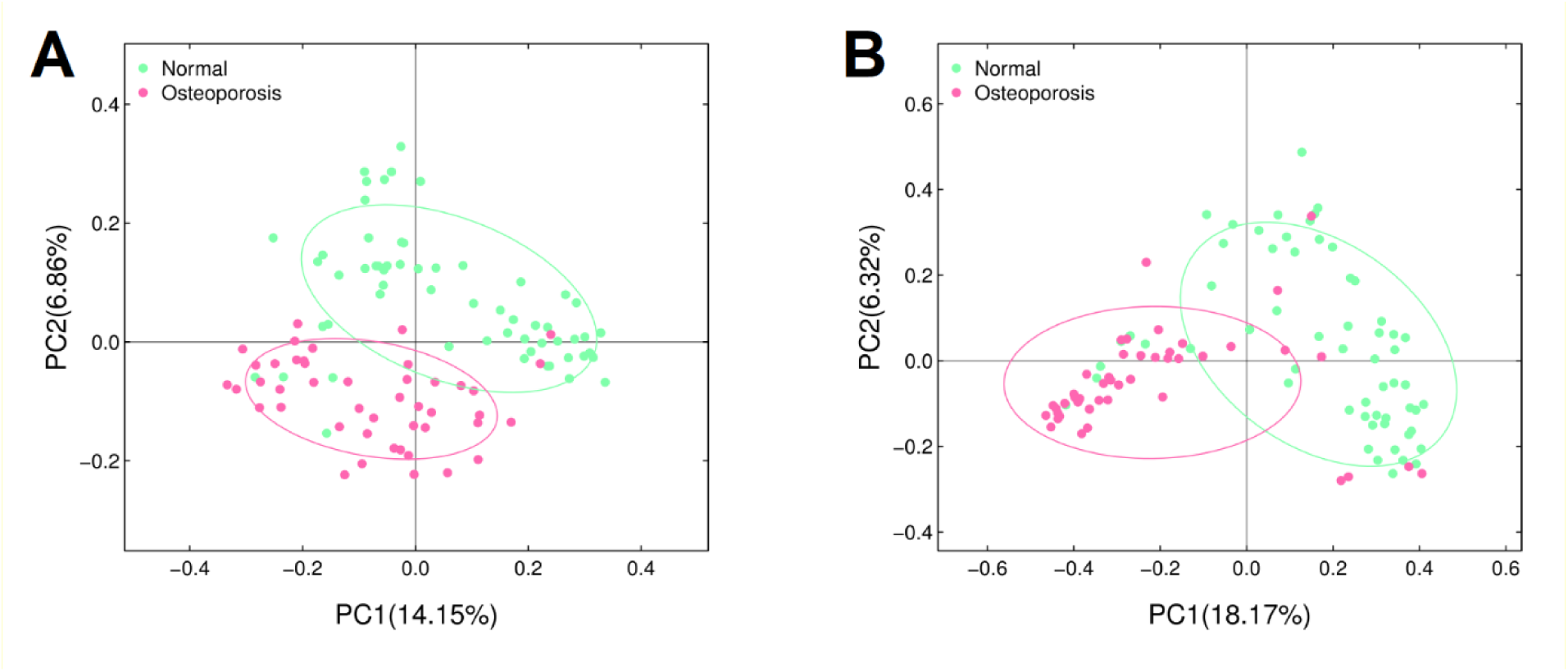
The normal subjects (green dots) and the osteoporosis cases (red dots) showed a separation in the principal coordinates analysis (PCoA) calculated with unweighted UniFrac distance (A) and Bray-Curtis dissimilarity (B).

### Differences in taxonomic abundance

We applied linear discriminant analysis effect size (LEfSe) software to explore the changes of the bacterial community in the osteoporosis all taxa level. Interestingly, a series of taxa exhibited differential abundance between two groups. At the genus level, the abundance of 29 genera (e.g., Roseburia and Bacteroides) was significantly higher in the osteoporotic cases than in the normal controls. On the other hand, the proportion of 15 genera (e.g. Streptococcus and Dorea) was decreased in the osteoporosis group (Figure 3 and Table 2).

**Figure 3.**
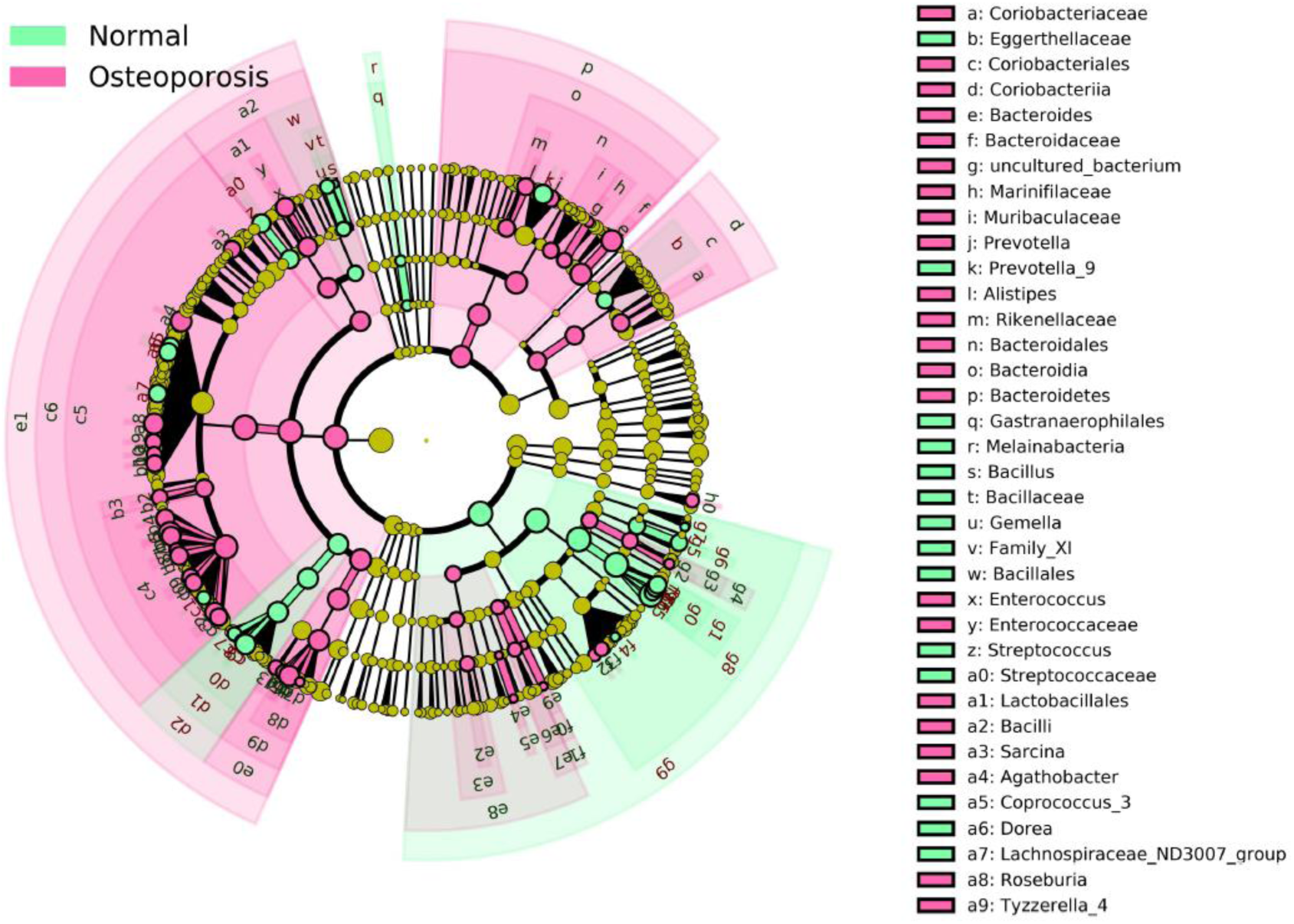
Taxonomic cladogram generated by LEfSe software showing alterations to gut bacterial communities in osteoporosis group.

To test whether the distinctive microbiota composition can classify osteoporotic status, we trained a random forest model with all study subjects using the profiles of genus. As indicated by the receiver operating characteristic (ROC) curve, the variable of genus abundance was effective to classify osteoporosis cases from controls (AUC=0.93, 95% CI 0.87-0.98, Figure 4A). A set of genera provided most of the discriminatory power, such as Roseburia, Streptococcus, Dorea and Flavonifractor (Figure 4B).

**Figure 4.**
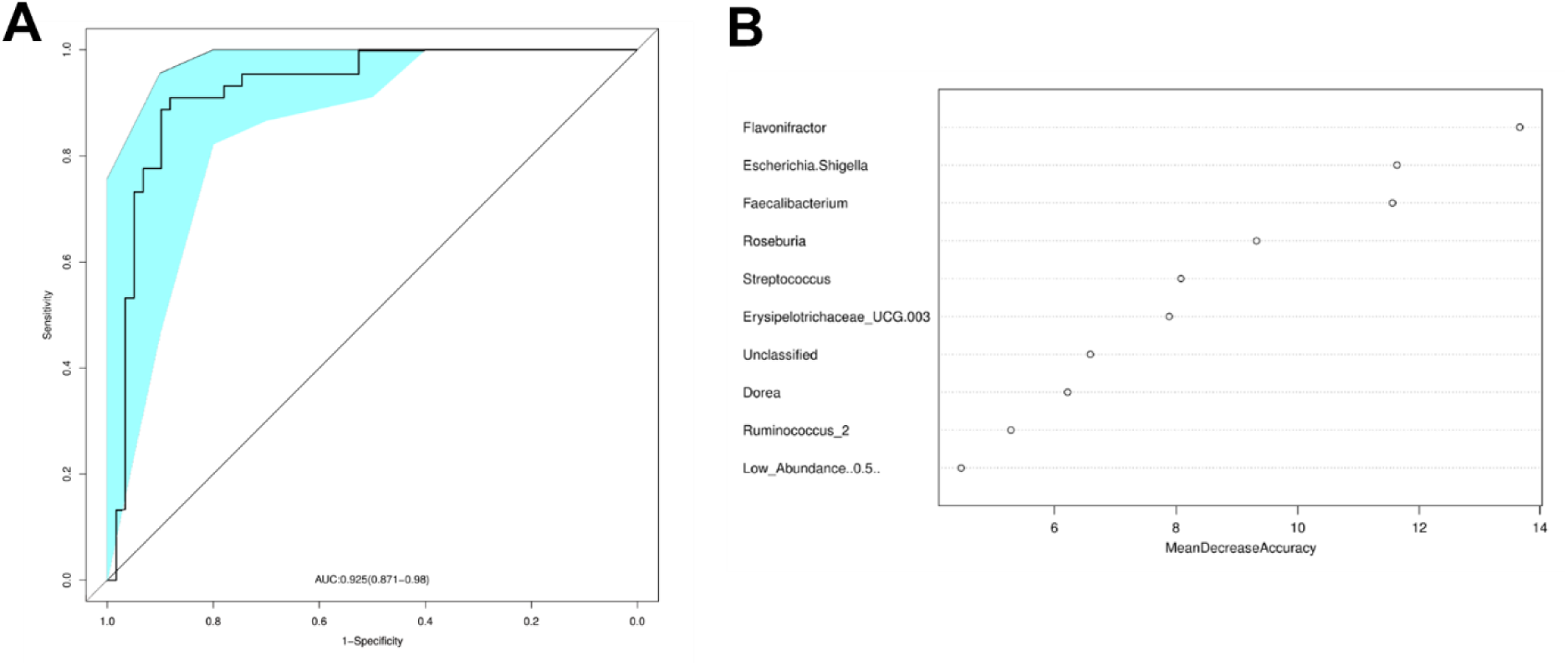
Gut microbiota signature which can be used to discriminate osteoporosis patients from normal controls. (A) Receiver operating characteristic curve for leave-one-out cross validation. (B) The top important genera in the discriminatory model as measured by mean decrease accuracy.

**Figure 5.**
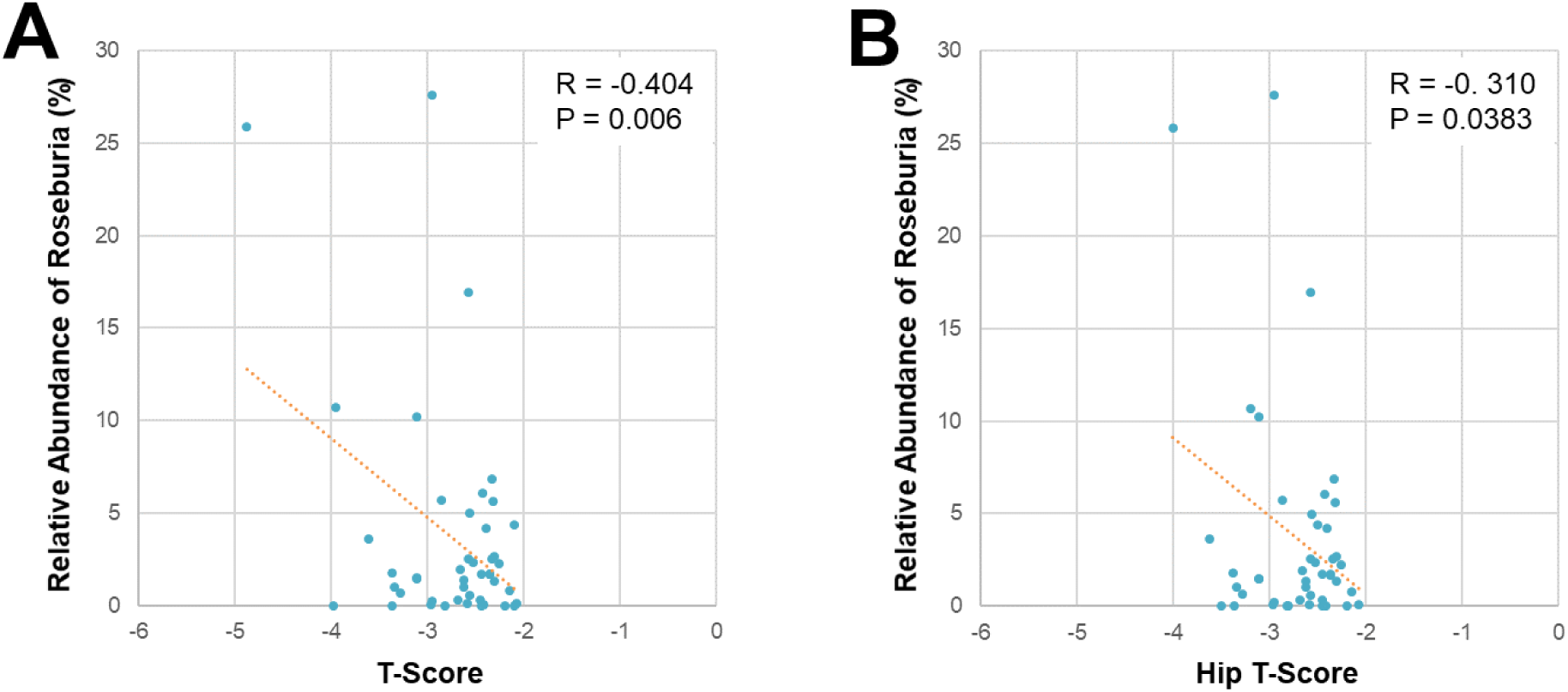
Correlation between bone mass level and the relative abundance of the genus Roseburia. (A) T-score. (B) Hip T-score.

### Gut microbiota and bone mass indicators

To evaluate whether osteoporosis leads to changes in the correlation between bone mass parameters and various taxa of the gut microbiota, we further performed Pearson’s correlation analysis. We identified only one genus, Roseburia, was negatively correlated with both T-score and Hip T-score (P < 0.05). In other words, the abundance of Roseburia genus was changed along with not only osteoporosis status but also severity of bone loss.

## Discussion

Conventional osteoporosis therapies focused on mitigating the loss of bone associated with decreases of sex steroids (16). With increasing knowledge about the pathology of osteoporosis, clinical research of new targets for therapeutic intervention is directed to gut microbiome associated to bone resorption and formation (17). In this study, we conducted 16S rRNA sequencing to compare the composition of gut microbiota between osteoporotic patients and control subjects.

Our results showed significant differences in the alpha diversity and the abundance of specific taxa in gut microbiome associated with osteoporosis. To date, there have been only several studies analyzing the alternations of gut microbiota in osteoporotic patients (9–11). However, the pooled female and male together or number limitation resulted inconsistent findings. Here we focused on the gut dysbiosis of female subjects with osteoporotic condition. One of our most notable performance was a clear distinction between osteoporotic cases and control subjects. Random forest model trained with microbiota composition data can effectively classify osteoporotic status, with the area under ROC curve reaching 0.93. Such improvement suggested that the risk and progress of female osteoporosis may be to a certain extent predicable by gut microbiota characteristics.

In our study, gut dysbiosis was characterized by a series of differential abundant genera in osteoporosis group. For example, we found an increased abundance of Bacteroides genus in osteoporosis group, which is consistent with previous report (11). Bacteroides is one of the most dominant genera in both osteoporotic individuals and normal subjects samples (10). As previously reported, Bacteroidetes might be involved in bone formation and bone resorption by deposition and hydrolysis of serine dipeptide lipids (18). Another noteworthy genus was Roseburia belonging to Firmicutes phylum. As an important producer of various short-chain fatty acids (SCFAs), Roseburia mediated the changes of IGF-1 expression and contributed to bone growth (19). Consistent to our finding, Roseburia was also found to be decreased in osteoporosis in the research with mixed female and male subject (11). More importantly, our results showed a significant enrichment of Roseburia among osteoporotic individuals, which served as one of the most efficient features in random forest model for discriminating patients from controls. Such inconsistency may be due to sex difference in pathogenesis of osteoporosis, since the interplay between microbiota and bone loss was dependent on sex steroid (20, 21). To explain the possible inconsistency, we will further investigate the role of sex steroid involvement in osteoporosis based on larger cohort and animal experiments.

Another major strength of our research was that we identified some clinical association in clinical parameters to support our hypothesis.It was firstly reported that there were abnormal levels of diamine oxidase, D-lactic acid and lipopolysaccharides (LPS) in osteoporotic patients. Circulating endotoxin, mainly composed by LPS, was secreted from intestine, and the intestinal permeability determined the secretory levels. Estrogen deficiency can influence intestinal epithelial permeability through mediating estrogen-associated pathways (22). All these findings together pointed to a possible theory that the estrogen level and the abundance of various bacterial taxa were both altered in osteoporosis status (23). The combined effects of estrogen deficiency and gut dysbiosis on intestinal permeability can mediate bone loss and osteoporosis via influencing the host metabolism, endocrine, and immunity (24), which might provide potential targets for the clinical treatment of post-menopausal osteoporosis.

On the other hand, several limitations of our results should be taken into consideration. based on the design of cross-sectional study, it was not possible to clarify the causal relationship between gut microbiota and osteoporosis. Since samples were collected after diagnosis, the changes of gut microbiota might be either a consequence of osteoporosis or the cause. Therefore, further study was required with larger sample size or well-controlled interventions on gut microbiota.

In summary, we described the altered profiles of gut microbiota in female osteoporotic patients, and provided evidence for the relationship between dysbiosis and osteoporosis. Our results pointed towards to early prevention and clinical management of osteoporosis by monitoring the homeostasis of gut microbiota.

## Acknowledgements

This study is supported by the Hong Kong General Research Fund. (201306151203), HKU Small Project Funding (201409176257), Science and technology program of medical and health of Shenzhen health committee of the NPC (SZFZ2017057), “Miao miao” cultivation program of Shenzhen hospital of southern medical university (2017MM01 and 2018MM11), Shenzhen Baoan district science and technology program basic research project (2017JD006 and 2017JD085), Key Research & Development Plan, Hunan, China(2016JC2071) and Project of Hunan Committee, Hunan, China(20201923). and the Research Fund for Lin He’s Academician Workstation of New Medicine and Clinical Translation.

---

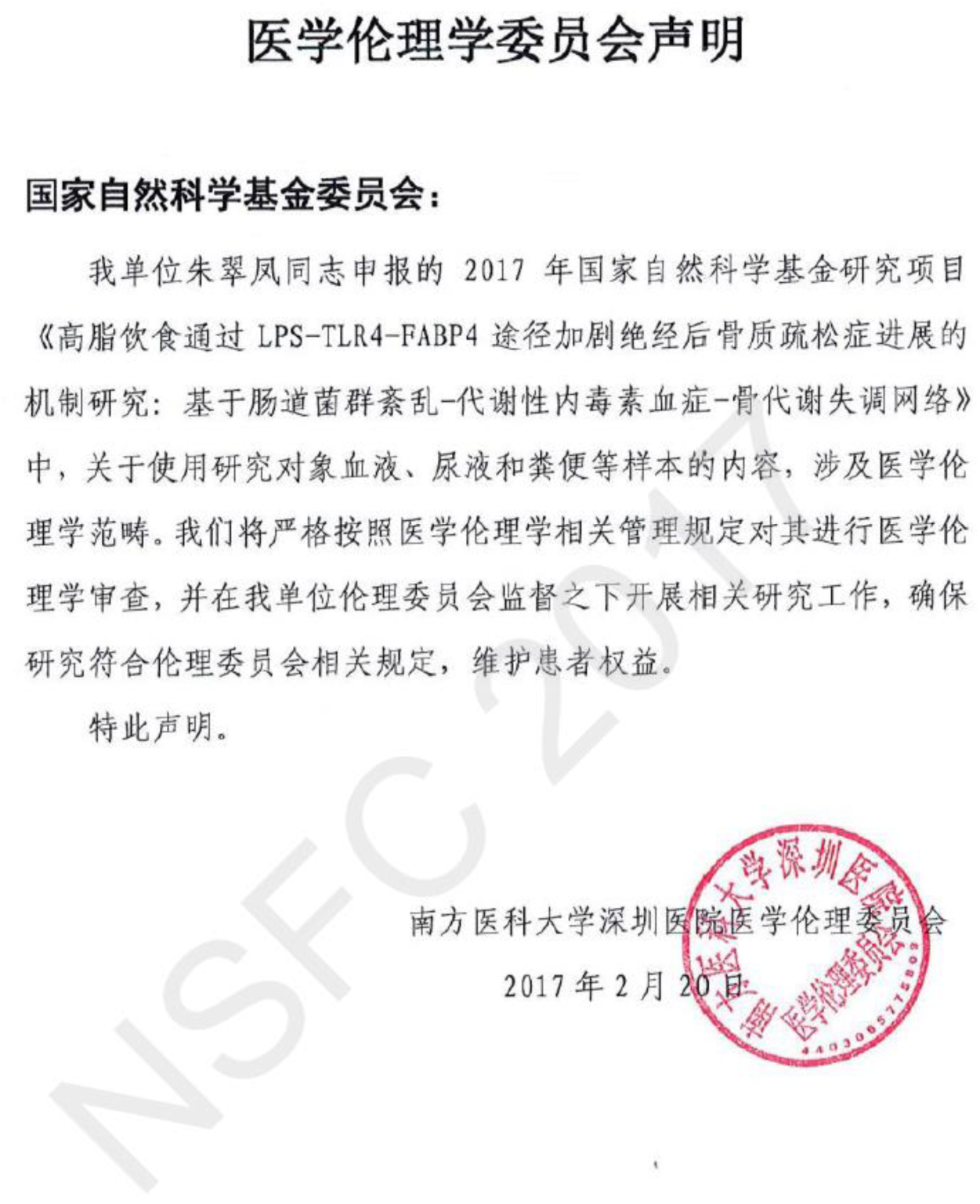

